# SQANTI-reads: a tool for the quality assessment of long read data in multi-sample lrRNA-seq experiments

**DOI:** 10.1101/2024.08.23.609463

**Authors:** Netanya Keil, Carolina Monzó, Lauren McIntyre, Ana Conesa

## Abstract

SQANTI-reads leverages SQANTI3, a tool for the analysis of the quality of transcript models, to develop a read-level quality control framework for replicated long-read RNA-seq experiments. The number and distribution of reads, as well as the number and distribution of unique junction chains (transcript splicing patterns), in SQANTI3 structural categories are informative of raw data quality. Multi-sample visualizations of QC metrics are presented by experimental design factors to identify outliers. We introduce new metrics for 1) the identification of potentially under-annotated genes and putative novel transcripts and for 2) quantifying variation in junction donors and acceptors. We applied SQANTI-reads to two different datasets, a *Drosophila* developmental experiment and a multi-platform dataset from the LRGASP project and demonstrate that the tool effectively reveals the impact of read coverage on data quality, and readily identifies strong and weak splicing sites. SQANTI-reads is open source and available for download at GitHub.

## INTRODUCTION

Short-read RNA sequencing (srRNA-seq) is the most common and cost-effective approach for studying the transcriptome. However, in srRNA-seq, transcripts must be inferred computationally, which can lead to inaccuracies in transcript identification (Liu et al. 2016; Newman et al. 2018). Recent advances in single-molecule long-read sequencing technologies have opened new avenues for transcriptome analysis (reviewed in (Marx 2023; van Dijk et al. 2023)). In long-read RNA sequencing (lrRNA-seq), full-length transcripts can be observed as single sequencing reads, allowing for direct transcript detection without the need for an assembly step. However, like any technology, lrRNA-seq is not without errors, and factors such as mRNA degradation, library preparation failures, and sequencing inacurracies can introduce biases into the data.

A database tracking bioinformatic tools for long-read sequencing (Amarasinghe et al. 2021) identifies numerous tools for the initial processing of lrRNA-seq data to assess the accuracy of base-calling and the length of the reads. These include pycoQC (Leger and Leonardi 2019), longQC (Fukasawa et al. 2020), nanoQC (De Coster et al. 2018), which offer a first-pass analysis of lrRNA-seq read quality. Other tools, such as SQANTI3 (Pardo-Palacios et al. 2024a), TALON (Wyman et al. 2020), FLAMES (Holmqvist et al. 2021), IsoSeq (https://isoseq.how/), and IsoTools (Lienhard et al. 2023), focus on evaluating transcript models inferred from the data. However, most current tools for lrRNA-seq read quality control were developed during the early stages of these technologies and are generally limited in the number of evaluated features and/or samples. As long-read sequencing technologies rapidly evolve, improving both in quality and experimental scope, the need for a comprehensive and comparative read quality assessment becomes increasingly critical.

Hence, the rapid decline in costs implies that the use of lrRNA-seq will continue to expand, with experimental designs involving multiple samples becoming more common (Glinos et al. 2022; Joglekar et al. 2024; Mahmoud et al. 2024; Patowary et al. 2024)). From a quality control perspective, this necessitates that datasets are homogeneous, without biases associated with experimental groups, and free of outliers. Moreover, the generated data must be sufficient to address the research questions that motivated the experiment. The increase in throughput now makes it possible to design experiments that include barcoding and multiplexing to balance library preparation and sequencing across experimental groups (Auer and Doerge 2010). This approach helps avoid confounding technical variation with the treatments of interest and facilitates discriminating between failed technical replicates and failed samples. Finally, technological advancements such as more accurate basecallers (https://github.com/nanoporetech/dorado) and the availability of novel library preparation methods such as MAS-Iso-Seq (Al’Khafaji et al. 2024), CapTrap (Carbonell-Sala et al. 2024), R2C2 (Volden et al. 2018), Nano3P-seq (Begik et al. 2023) or FLAM-seq (Legnini et al. 2019) motivates the need for tools that can easily evaluate how these improvements impact various aspects of the long-read data quality.

In this context, we present SQANTI-reads, an extension of SQANTI3 (Pardo-Palacios et al. 2024a), a tool originally designed for transcript model quality control, to jointly provide quality control metrics for long-read data and to analyze multiple samples for consistency and bias. We demonstrate that SQANTI3’s structural categories and other quality control metrics, repurposed in SQANTI-reads, are highly effective for assessing the homogeneity in a lrRNA-seq multi-sample experiment, identifying read quality control failures, and detecting outliers. Additionally, we have add new metrics that provide insights into the potential utility and discovery power of the data, including variation at donor/acceptor sites and identification of potentially under-annotated genes and mis-annotated transcripts. SQANTI-reads offers an extensive array of summary output tables, is customizable to accommodate any experimental design, and is available as an open-source, freely accessible tool.

## RESULTS

### SQANTI-reads can be used to evaluate long-read technology improvements

Long-read methods are rapidly improving and both Nanopore and PacBio are updating their instruments, kits and algorithms. Tools that can readily evaluate the broad impact on data quality of these technical improvements are highly needed to make informed decisions for downstream analyses. Our *Drosophila* dataset contained the same set of samples processed on the MinION and Promethion Instruments through several runs and analyzed with two base callers, the default Guppy and the new Dorado algorithm. A first question is whether Dorado effectively improved data quality without introducing biases and if data from several sequencing experiments could be merged. SQANTI-reads was used to address these questions. We first compared the Guppy and Dorado basecallers using SQANTI-reads metrics. As anticipated, Dorado resulted in more reads with assignable barcodes, a higher number of mapped reads, more reads aligning to annotated genes, more reads aligning to annotated transcripts, and more longer reads, without an increase in the proportion of reads with technical artifacts (Supplementary Figure 1, Supplementary file 2). This confirms that Dorado improves base-calling accuracy without introducing unwanted biases and motivated the selection of Dorado base-called reads for subsequent analyses.

**Figure 1:**
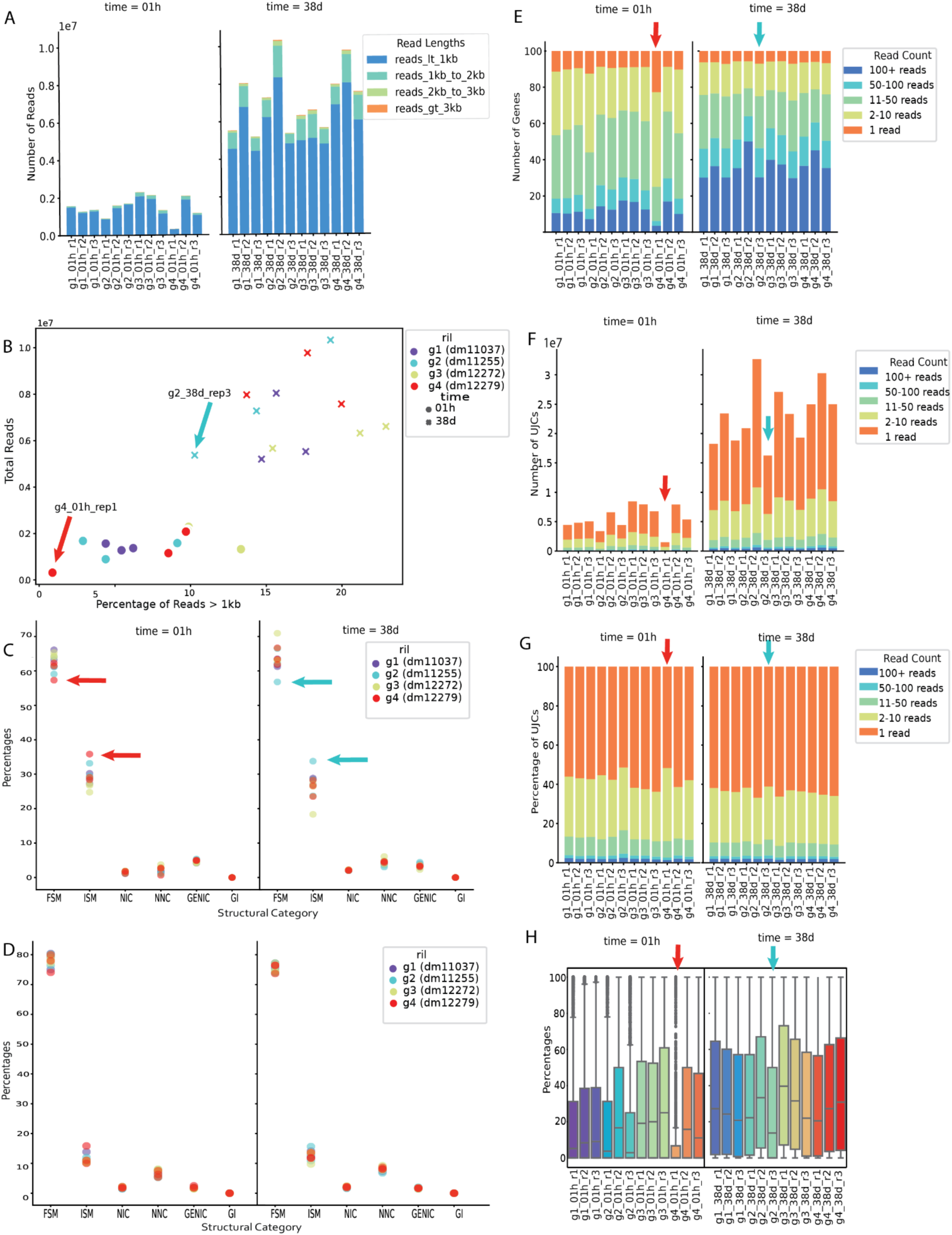
SQANTI-reads analysis of *Drosophila* samples. A) Number of mapped reads by experimental group labeled with read length. B) Percentage of mapped reads >1kb vs percentage of reads that are FSM. Dots represent early stage (0-1 hours after enclosure) and crosses indicate adult stage (3 to 8 days old). The four genotypes are indicated with four different colors. C) Percentage of reads mapping to genes in each SQANTI3 structural category D) Percentage of reads mapping to genes in each SQANTI3-QC structural category for reads >1kb E) Number of genes detected with breakdown by the number of reads mapped to each gene. F) Number of UJCs detected with breakdown by the number of reads associated with each UJC. G) Proportion of UJCs detected with breakdown by the number of reads associated with each UJC. H) Distribution of the percentage of FSM reads by gene across samples.

In the *Drosophila* experiment, libraries were barcoded, pooled, and multiplexed across different MinION and PromethION runs, with a re-pooling step between the two machines. We used SQANTI-reads to compare the quality of the MinION and the PromethION runs, and to evaluate the consistency of the MinION and PromethION technology across technical replicates. The first MinION run (TR1) had higher percentages of reads with NIC/NNC and with non-canonical junctions compared to the second and third technical MinION runs (TR2, TR3) and compared to the PromethION runs for the same libraries (Supplementary Figure 2). MinION runs TR2 and TR3 were similar in their quality metrics (described below) to the PromethION run of the same samples, and the technical replicates on the PromethION were similar. These results indicate that the technology performs consistently across instruments and runs. Based on these SQANTI-reads QC results we aggregated data across technical replicates to further evaluate the quality of the lrRNA-seq experiment. Although TR1 had lower quality than the other technical replicates, the overall read numbers were low and we decided to keep this technical replicate in our evaluation of the samples.

### SQANTI-reads metrics can be used to evaluate the global quality of the lrRNA-seq experiment

In a multi-sample lrRNAseq experiment, all samples should be of similar quality. SQANTI3 uses the FSM structural category to identify long-read sequences whose junctions are consistent with an annotated transcript model. However, for a lrRNA-seq experiment to accurately reflect the analyzed transcriptome, the reads should ideally also capture the distribution of transcript lengths of the expressed transcriptome. The distribution of transcript lengths depends on the species with *Drosophila* having overall less complex and shorter transcripts than human (Supplementary Figure 3). A dataset with reads substantially shorter than the targeted transcriptome but with still a high number of FSM indicates capture of short transcripts, while combining shorter than expected reads with a high proportion of ISM may indicate RNA degradation. We looked at these values for an initial assessment of the quality of the *Drosophila* experiment.

First, we compared the number of reads and length distributions for all samples. The difference in sequencing depth between the two developmental stages was evident, and motivated by the the additional PromethION run on the 3-8 day samples (Figure 1A). For all samples, most reads were shorter than 1kb with less than 20% of them above the 1 kb threshold (Figure 1B, Supplementary Figure 4A). While between 53% and 67% of the reads across samples were classified as FSM, 20% to 38% were labeled as ISM, and NIC/NNC were under 10% of the reads (Figure 1C). For the reads greater than 1 kb, between 73% and 82% of the reads were FSM while 10% and 18% were ISM (Figure 1D). These results suggest that the *Drosophila* dataset contains a significant proportion of short reads leading to incomplete transcript sequences.

To further understand how read quality affects gene and transcript quantification, we examined these metrics aggregated by gene and UJC. We found that, despite the large sequencing depth differences between developmental stages, the number of detected genes was only slightly lower in the 0-1 h samples (Figure 1E). However, these genes were quantified with fewer reads (80% genes with < 50 reads) than the 3-8 d samples, which had between 30% and 50% of genes with more than 100 reads (Figure 1E). Interestingly, when evaluating UJC we found that, while the number of UJC mirrored the sequencing depth pattern (Figure 1F), with 3-8 d samples showing five times more UJC than 1 h samples, and a larger number of FSM and ISM UJC, there were many additional UJC detected by fewer than 10 reads, and usually by a single read (Figure 1F & 1G) and these UJC were most frequently NIC/NNC (Supplementary Figure 4B & 4C). Downstream analyses would therefore need to address whether this represents novel low-expressed transcripts or technology errors. In contrast, the percentage of FSM reads between the two time points differed by less than 1x in all replicates (Figure 1H). These results indicate that the higher sequencing depth of the 3-8 d samples does not change the number of detected genes or annotated transcripts (FSM). The higher read depth per gene/ UJC suggests that more genes and transcripts will be able to be quantitatively evaluated in the 3-8 day samples compared to the 0-1 hour samples.

In the *Drosophila* data, we noticed two samples (RIL 12279 rep 1 0-1 hour red arrow; RIL 11255 3-8 day rep 3 - teal arrow) that had the lowest percentage of FSM and highest percentage of ISM in the 0-1 hour and 3-8 day groups respectively (Figure 1C). To determine whether these two samples were of overall lower quality than the rest, we examined their SQANTI-reads metrics. We found that RIL 12279 rep 1 had a lower proportion of FSM across all genes (Figure 1H) and a higher proportion of genes quantified with only one gene (Figure 1F), while RIL 11255 rep 3 had a similar gene (Figure 1H), UJC (Figure 1G) and % FSM in genes (Figure 1H) than other 3-8 day samples. We concluded that RIL 12279 rep 1 0-1 hour is a low-quality sample.

Altogether, this example shows that SQANTI-reads metrics can be used to compare samples and experimental conditions in a multi-sample experiment, detect outliers, and suggest points of attention for downstream data processing.

### SQANTI-reads metrics can be used to identify systematic differences among samples

The previous example demonstrated that SQANTI-reads metrics are effective in assessing dataset consistency. However, SQANTI-reads evaluates over 35 quality metrics, making it challenging to determine which features contribute to potential differences among samples. We include Principal Component Analysis (PCA) analysis to identify which metrics are the most relevant for quality variability when there are differences among samples or between groups. The percentage of reads and UJCs in each structural category, percentage of artifact reads (RT-switching, non-canonical junctions and intrapriming), percentage of junctions in each category, as well as length metrics, are included in the PCA.

We applied SQANTI-reads PCA analysis of quality features to investigate differences in read quality among various long-read sequencing methods used in the LRGASP challenge (Pardo-Palacios et al. 2024b), focusing on the WTC11 dataset. The analysis revealed that WTC11 samples clustered based on the long-read technology applied (Figure 2A). Specifically, PC1, which explains 56% of the variance, distinguished cDNA ONT samples from those generated by the other two technologies, while PC2, accounting for 35% of the variance, highlighted differences between dRNA ONT and cDNA PacBio. To further explore these differences, we examined the loadings for each principal component. Quality features with the highest positive loadings in PC1 included the number of reads, the percentage of reads and the proportion of UJCs in the NNC category, while features with high negative loadings included Intergenic and Genic Genomic reads. Several junction-related variables also exhibited high absolute loadings on PC1 (Figure 2B). SQANTI-reads plots confirmed these structural category differences between cDNA ONT samples and other library preparations. cDNA ONT had both the highest proportion of NNC reads and UJCs, (Figure 2C and 2D) and also had the lowest proportion of intergenic reads (Figure 2C). Other differences in sequencing throughput and junction characteristics were also confirmed (Supplementary Figure 5).

**Figure 2:**
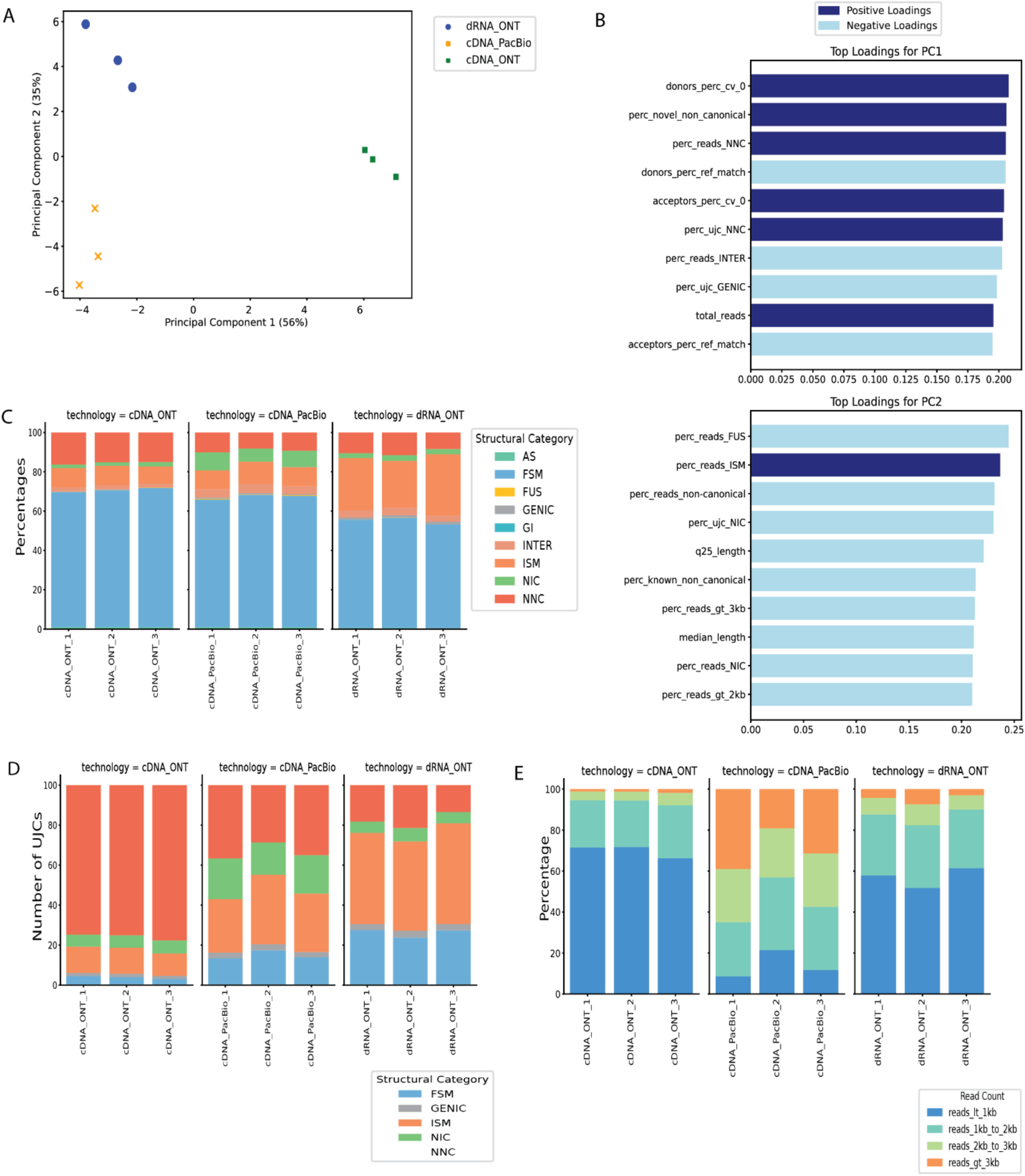
SQANTI-reads PCA analysis of LRGASP WTC11 samples. A) PCA using SQANTI-reads quality features. B) Top 10 Loadings for PC1 and PC2. C) Distribution of reads in SQANTI structural categories. D) Distribution of UJCs in structural categories E) Distribution of read lengths for all mapped reads.

Upon examining the feature loadings for PC2, we found that variables with high contributions included several metrics related to read length (Figure 2B). Consequently, we evaluated the SQANTI-reads “Lengths of All Mapped Reads” plot for this experiment. Indeed, we observed that the cDNA PacBio method produced a significantly higher proportion of reads between 1-2 kb, 2-3 kb, and greater than 3 kb, as suggested by their negative loadings, compared to the dDNA ONT method, which predominantly generated reads shorter than 1 kb (Figure 2E). Similarly, percentage of reads assigned as ISM with high positive values was higher in dRNA ONT samples (Figure 2 C).

In conclusion, we showed that the SQANTI-reads PCA analysis is an effective tool for uncovering significant read quality differences between long-read sequencing methods and for revealing unexpected technology biases.

### SQANTI-reads identifies potentially under annotated genes

Long-read data often reveal a large number of sequences that cannot be exactly matched to existing annotations. In many cases, these UJC belong to annotated genes and are identified by only a few reads, as illustrated by the SQANTI-reads analysis in Figures 1 and 2. This suggests they could be either low-expressed transcripts or technological artifacts. However, in some instances, a high proportion of reads in a gene may correspond to the same novel UJC, indicating the possibility of a previously unannotated transcript that warrants closer examination. SQANTI-reads includes a customizable decision tree to identify such cases (see Methods, Figure 3A). Basically, the tool identifies genes with a high (R > threshold) number of reads, with novel UJC containing a large fraction (Q > threshold) of the genés splice sites and capturing a high proportion (P > threshold) of the reads. We applied this approach to the WTC11 PacBio data using default parameters (R=100, P=20 and Q=80). The logic for defining expressed genes as well annotated or underannotated is described in Figure 3A. The annotation category for all expressed genes (default: number of reads in gene (R) > 100) are provided in the gene_classification.csv file. From the set of expressed genes, we identified 8,556 well annotated genes, 88% of which have a well covered annotated transcript (>20% of total gene coverage) (Figure 3B). We also identified 101 genes, for which there are no reads with an FSM match to an annotated transcript. Of these 54% have a well covered UJC. For all expressed genes with well covered unannotated UJC, we identified 424 that contained most of the observed junctions in that gene (>80%), and we label these putative novel transcripts (Figure 3C). Of these, 316 were NIC and 108 were NNC. The SQANTI-reads output for putative novel transcripts is included in the putative_underannotation.csv file (Table 2).

**Figure 3:**
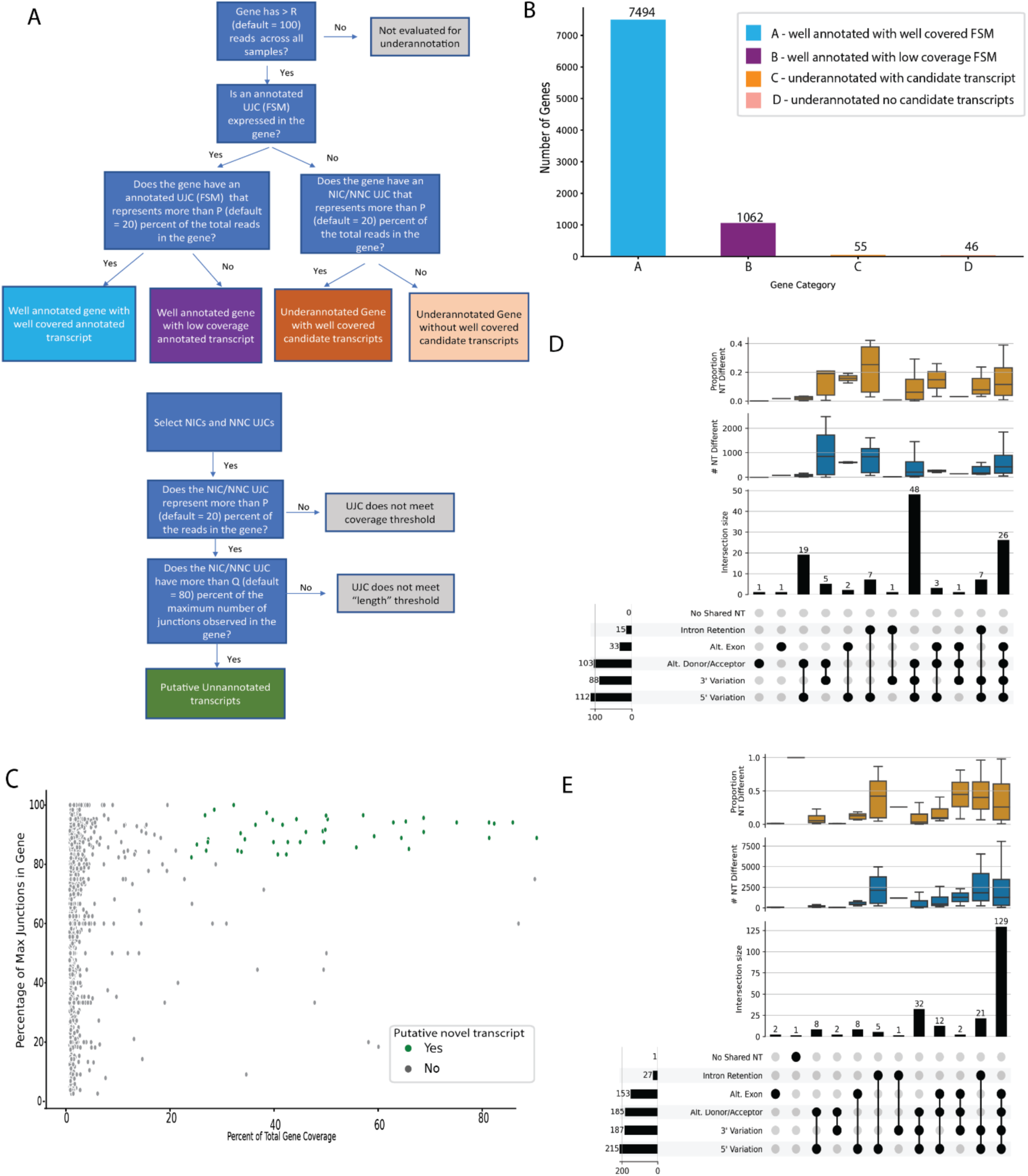
Evaluation of WTC11 PacBio samples for under-annotated genes. A) Decision tree for classifying genes as well annotated or under-annotated and classifying transcripts as putative novel candidate transcripts. B) Number of genes by their annotation status according to SQANTI-reads parameters. C) The coverage (percent of total reads) vs length (percentage of maximum junctions) for all UJCs in underannotated genes with well covered candidate transcripts. UJCs that meet the thresholds for putative novel transcripts are colored in green. D&E) Comparison of putative candidate novel transcripts with the most expressed annotated transcript in that gene for well-annotated genes with well-covered FSMs (D) and with an FSM detected but without < 20% of the total reads in the gene(E).

For genes with at least one putative novel transcript (R>100, P>20, Q>80) and an annotated transcript (FSM) we selected the FSM with the highest proportion of reads. We then compared the structure of the putative novel transcripts to the most expressed annotated transcript using TranD (Nanni et al. 2024). For genes with an annotated transcript that is relatively highly expressed (>20% of the reads for that gene), 103 putative candidate transcripts differed from the annotated transcript by donor/acceptor variation, suggesting a possible alternative splice site. In addition, 10 putative candidate transcripts had an extra exon; 15 a skipped exon and 9 with both missing and skipped exons relative to the most expressed annotated UJC (Figure 3D). For the genes where the annotated transcript represented less than 20% of the reads in that gene, the putative novel transcript differed from the annotated transcript by an alternative exon in 147 cases (33 extra exons, 86 skipped exons, and 43 with both an extra and skipped exon) (Figure 3E). Details and scripts for this analysis are provided in the Supplementary Methods.

This analysis shows that SQANTI-reads can readily identify under-annotated genes and flag putative novel transcript models that contain interpretable alternative exonic patterns that deserve further attention.

### SQANTI-reads metrics for donors/acceptors identify noisy splicing and potentially novel splice-sites

SQANTI-reads calculates the mean, standard deviation and coefficient of variation (CV) for all expressed annotated donors/acceptors (Figure 4A). A value CV > 0 indicates variability in the donor/acceptor, with higher CV values indicating more variability. Variability around a splice-junction may be due to weak splicing (Wang and Marín 2006) or to technology errors and mapping accuracy, for example due to junction ambiguity (Li 2018). We evaluated these metrics on the WTC11 dataset for reference junctions with at least 10 reads. We found similar patterns in the variability (CV > 0) in donors and acceptors (Figure 4B). All three technologies identify donors and acceptors with variability around the splice site (CV > 0). Donors/acceptors with CV > 0 consistently across the three technologies, are highly suspicious of ‘noisy’ splicing or a weak splice site and may be worth follow up (Supplementary Figure 6A and 6B). The SQANTI-reads output file cv.csv identifies the donors/acceptors with CV > 0 making it straightforward to follow up on particular locations with tools such as Integrative Genomics Viewer (IGV) (Robinson et al. 2011).

**Figure 4.**
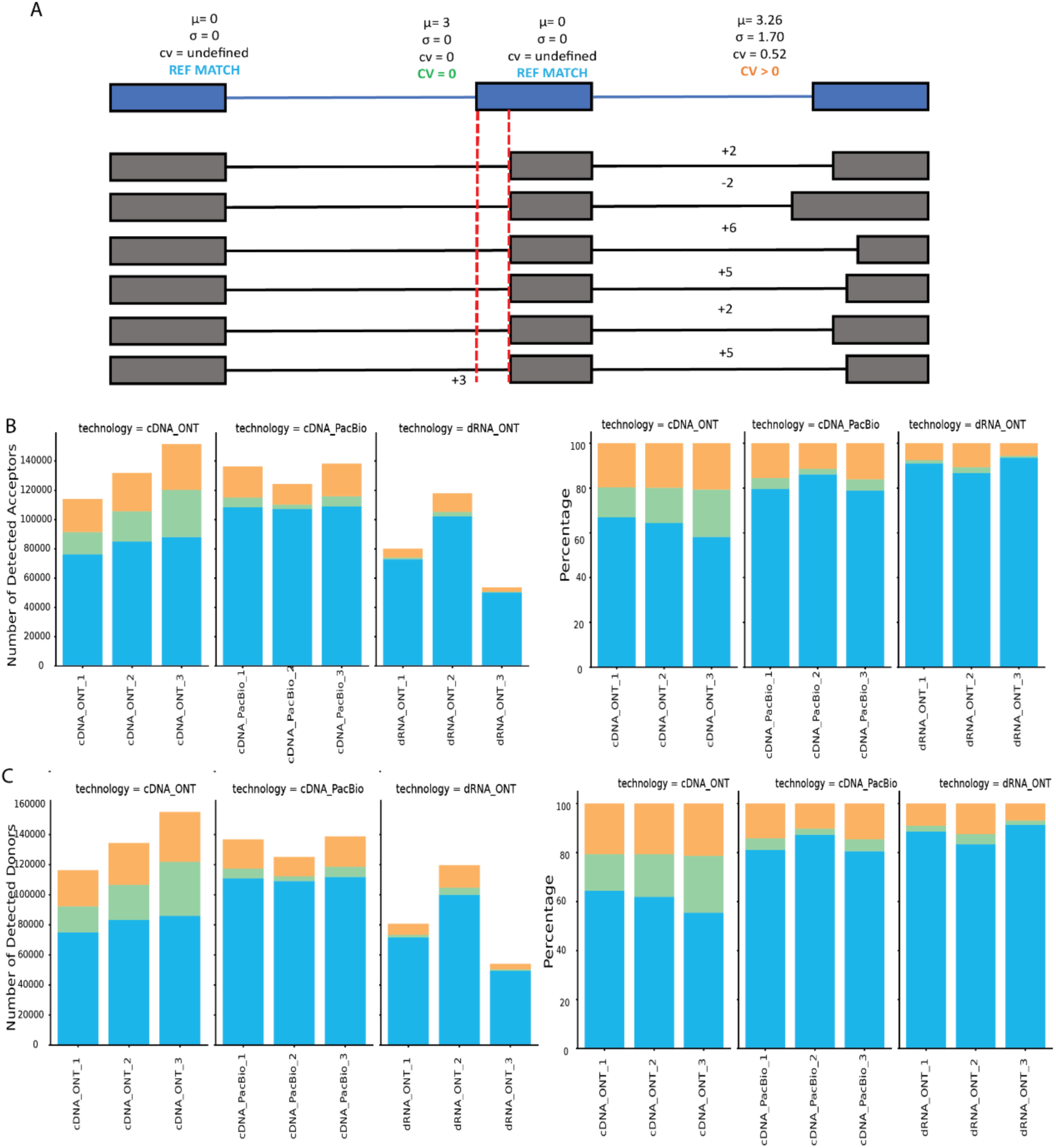
Variation in donors/acceptors. Metrics are only calculated for annotated donor/acceptors with a minimum threshold of reads (10 by default). A) Metrics for the classification of donor/acceptor variation. When all reads align to the annotated donor/acceptor this is classified as a reference match (Ref Match). When all reads align to the same donor/acceptor location, but this is not the annotated position this is classified as CV = 0. When reads align in multiple positions in proximity to an annotated donor/acceptor this is classified as CV > 0. B) Classification of the number (left) and proportion (right) of detected acceptors faceted by technology. C) Classification of the number (left) and proportion (right) of detected donors faceted by technology.

A reference match junction indicates that the splice signal is strong (Wang and Marín 2006; Dent et al. 2021). Results for cDNA PacBio and dRNA-seq were similar, with both showing a higher number and proportion of reference match donors/acceptors compared to cDNA ONT (Figure 4B and C) despite these technologies detecting similar numbers of FSM UJCs (Figure 2D). We compared the FSM UJCs identified by the three methods (Supplementary Figure 7A). Most of the FSM UJCs were detected by all three technologies (n = 19690), with a similar number detected by cDNA PacBio only (n = 8110) and cDNA ONT (n = 9360) only. We hypothesized that the difference in the number of reference match donors/acceptors was potentially due to longer transcripts with more junctions being detected in cDNA PacBio compared to shorter transcripts with fewer junctions in dRNA ONT and cDNA ONT. For the FSMs detected only in one technology we plotted the distribution of the number of junctions and confirmed that the cDNA PacBio FSM transcripts had a larger number of junctions compared to dRNA ONT and cDNA ONT (Supplementary Figure 8). This agrees with cDNA PacBio showing longer reads than both ONT technologies in the WTC11 dataset (Figure 2E).

Splice junctions may differ from annotated sites due to the presence of novel donors/acceptors. The category CV = 0 identifies donor/acceptors with no variability but differing from the annotated donor/acceptor, representing strong candidates for *bona fide* alternative splice sites. We evaluated the donors and acceptors with CV = 0 in all the technologies (Supplementary Figure 6E and 6F). We identified 51 donors and 34 acceptors with CV=0 in all three technologies. Of these, 76% of donors and 65% of acceptors are within 12nt of an annotated donor/acceptor indicating a potential mis-annotation of the splice-site (Supplementary Figure 9). These donors and acceptors with CV = 0 across all technologies were detected by a minimum of 10 reads and in some cases could be detected with > 100 reads (Supplementary Figure 10). Supplementary Figure 11 shows an example of one of the reference donors with CV = 0 detected across all 3 technologies. This donor is in the H2AZ1 gene at position 99,948,814 on chromosome 4. All the reads associated with this donor map to position 99,948,811 which is 3 nts away from the annotated donor position (Supplementary Figure 11). Donors and acceptors with CV = 0 detected consistently across samples and/or technologies indicate potential robust detection of alternative splice sites.

## DISCUSSION

The increase in throughput and generalization of lrRNA-seq has led to the growing popularity of experiments containing many samples. Multiple lrRNA-seq studies of lrRNA-seq have shown that, despite technological improvements, biases associated with read length and accuracy are still present in lrRNA-seq data (Amarasinghe et al. 2020; Delahaye and Nicolas 2021). These biases vary between the major LRS platforms and with the introduction of new instruments and chemistries. Other types of systematic biases, such as those introduced by batches utilized in large experiments, may also occur. Benchmarking studies have also revealed that transcript reconstruction methods can yield highly different results due to the varying strategies used to resolve data inaccuracies (Pardo-Palacios et al., 2024b). This highlights the need for tools that can comprehensively evaluate the characteristics of raw long-read data before making decisions on downstream analyses. Direct examination of read quality enables the researcher to evaluate the experiment for consistency and identify any outlier samples and any systematic differences in read quality between sample groups. We have developed SQANTI-reads as a tool that enables a comprehensive assessment of reads obtained from a LRS experiment. We leverage the widely adopted SQANTI3-QC framework and we add several metrics designed specifically for the assessment of reads to provide an effective strategy for evaluating the quality of lrRNA-seq multi-sample experiments. Mapped reads are summarized by their inferred junctions, start and end positions and using SQANTI3 categories and sub-categories each read is classified. A summary of all the observed patterns of junctions, and the distance between annotated and observed donors and acceptors are tallied by sample. If meta-data is included in the design file, sample groupings can be used to compare SQANTI-classifications and junction metrics. Alternate groupings based on different columns in the meta-data are enabled without needing to rebuild the classification and junction files.

For example, in Oxford Nanopore Technology (ONT) the existence of multiple platforms at different price points for different numbers of pores (Flongle, MinION, GridION, PromethION) but with the same library protocols means that in a large experiment, samples can be initially evaluated at low cost on one of the lower throughput platforms (Flongle MinION) and if samples are of sufficient quality, then can be run on higher throughput platforms (GridION, PromethION). Sample multiplexing and running on multiple ‘lanes’ is good experimental design practice (Auer and Doerge 2010). Our SQANTI-reads analysis of the Drosphila dataset illustrates such scenario. In this case the pool for the 0-1 hour samples was initially evaluated on the MinION. The resulting data from TR1 were unbalanced and had relatively short reads with a high proportion of ISM. These observations enabled adjustment of the library concentrations and run parameters, and a second minion run resulted in fewer ISM and more balance in read numbers across libraries. A PromethION run based on the rebalanced libraries efficiently used the sequencing resources across samples. The same procedure was deployed with the 3-8 day samples, and while rebalancing helped ensure efficiency of the subsequent two PromethION runs. This example illustrates both good experimental design practices for large experiments and the utility of SQANTI-reads in assessing data quality at early stages of data acquisition, enabling corrections that lead to a successful sequencing experiment.

Another important aspect of lrRNA-seq multi-sample experiments is the ability to quickly assess whether the data can address the biological questions that motivated the study. This includes, among other things, whether genes and transcripts are sufficiently quantified and if potential novel transcripts are adequately supported. While these questions may ultimately be answered after full data processing with transcript reconstruction algorithms, users may find it helpful to evaluate support directly from the raw data.

SQANTI-reads provides information on the distribution of reads across genes and UJC (a raw-data proxy for transcripts) and introduces new metrics for identifying variations in donors/acceptors, under-annotated genes, and putative novel transcripts for further evaluation. These metrics enable the researcher to quickly determine if more reads are needed, and whether there are highly expressed putative novel transcripts potentially worth detailed experimentation.

The examples presented in this work demonstrate that SQANTI-reads is flexible and customizable, allowing users to explore the impact of various experimental design factors on read, UJC, and donor/acceptor properties, as well as identifying potential novel transcripts. The output from SQANTI-reads can be easily mined for additional insights and used to direct attention and resources toward interesting and novel features of lrRNA-seq experiments. We expect SQANTI-reads to become an essential tool for the QC of multi-sample lrRNA-seq datasets.

## METHODS

### SQANTI-reads basics

SQANTI-reads is an adaptation of SQANTI3 designed to evaluate individual reads rather than transcript models. It allows for the comparison of multiple samples, providing quality control results across the entire experiment. Several new features have been introduced to address the specific needs of QC in multi-sample experiments, while some functionalities of SQANTI3 have been removed as they are not applicable to read-level processing. Table 1 highlights the major differences between SQANTI3 and SQANTI-reads, emphasizing the new features of SQANTI-reads, while Table 2 lists the names and descriptions of the output files.

**Table 1.**
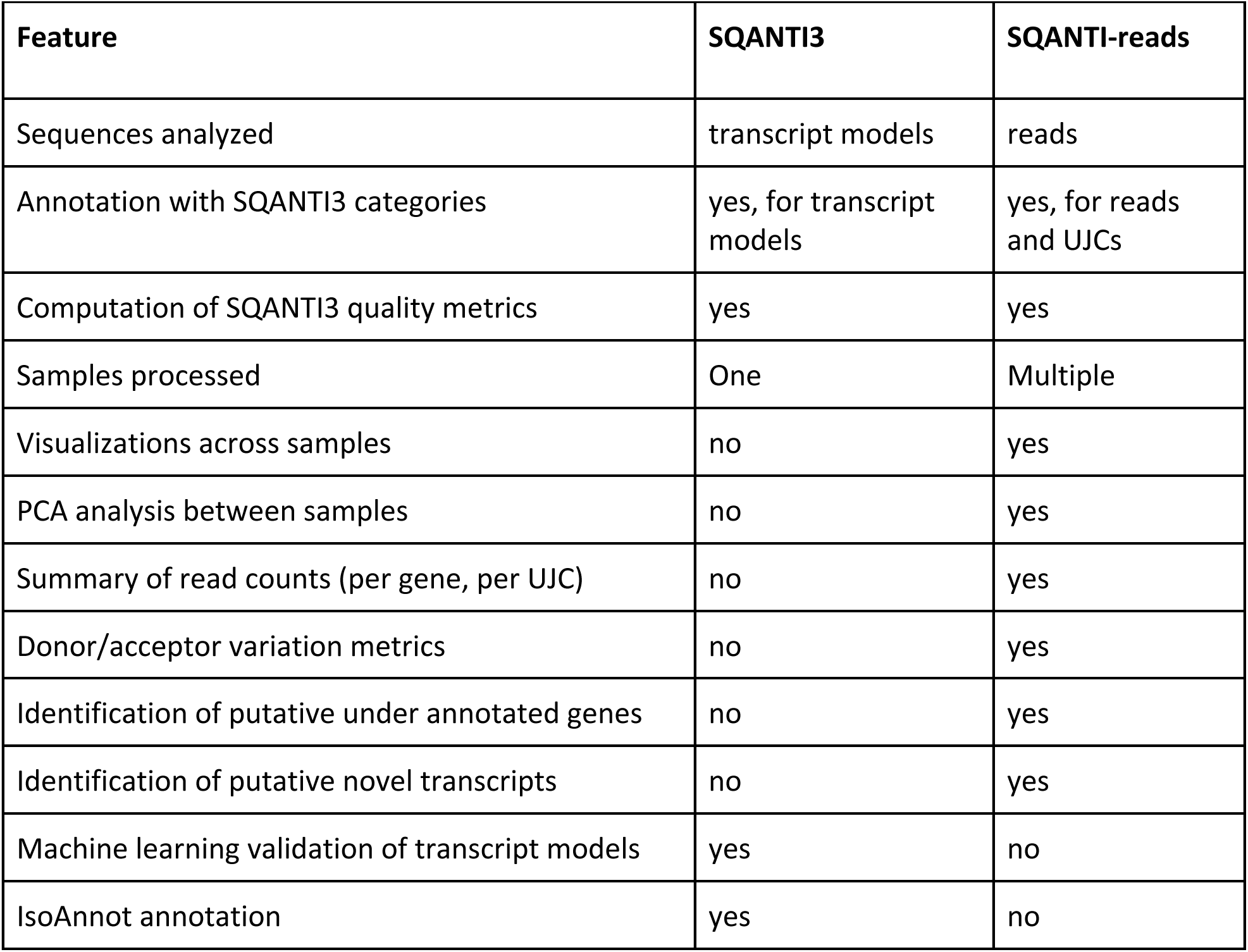
Comparison between SQANTI3 and SQANTI-reads.

**Table 2.**
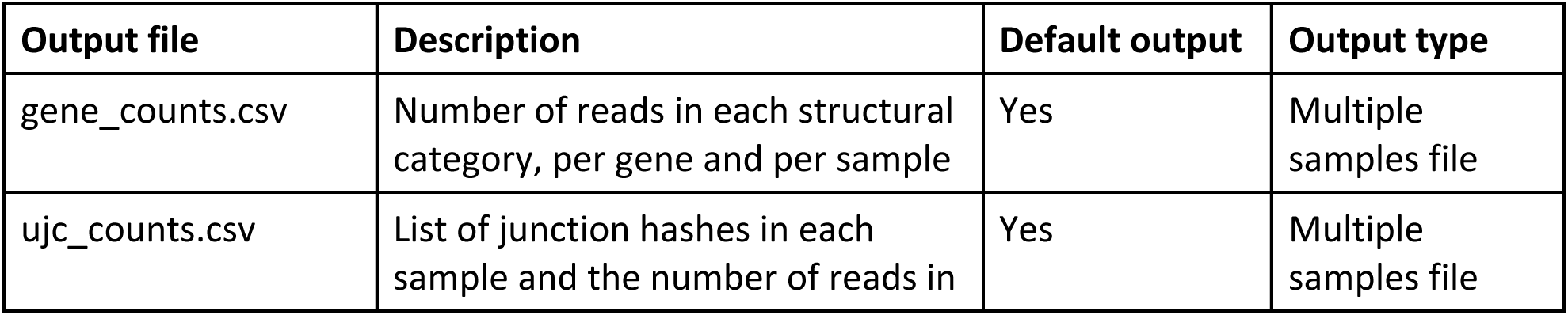
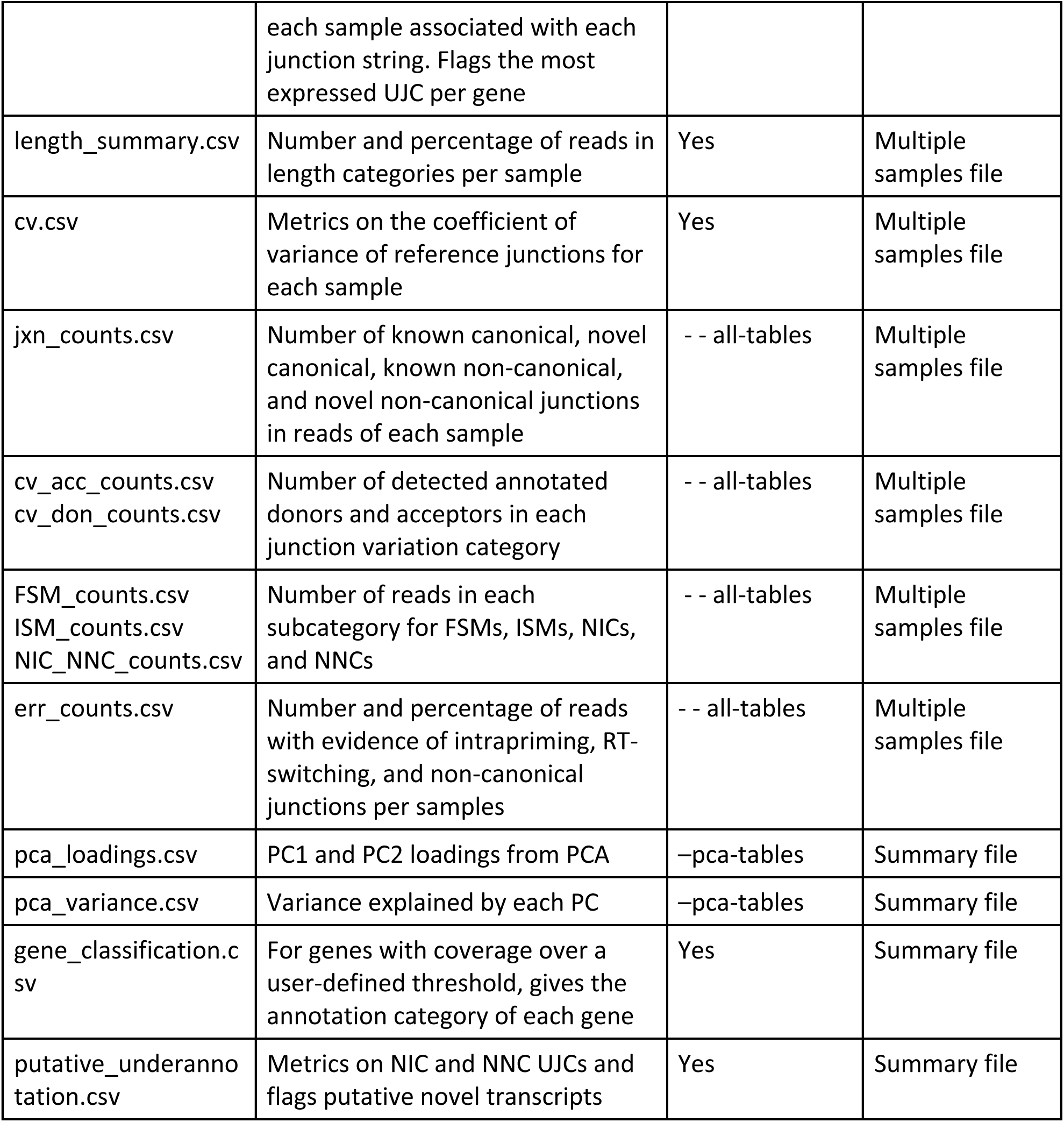
SQANTI-reads specific output files.

| Output file | Description | Default output | Output type |
| --- | --- | --- | --- |
| gene_counts.csv | Number of reads in each structural category, per gene and per sample | Yes | Multiple samples file |
| ujc_counts.csv | List of junction hashes in each sample and the number of reads in each sample associated with each junction string. Flags the most expressed UJC per gene | Yes | Multiple samples file |
| length_summary.csv | Number and percentage of reads in length categories per sample | Yes | Multiple samples file |
| cv.csv | Metrics on the coefficient of variance of reference junctions for each sample | Yes | Multiple samples file |
| jxn_counts.csv | Number of known canonical, novel canonical, known non-canonical, and novel non-canonical junctions in reads of each sample | - - all-tables | Multiple samples file |
| cv_acc_counts.csv<br>cv_don_counts.csv | Number of detected annotated donors and acceptors in each junction variation category | - - all-tables | Multiple samples file |
| FSM_counts.csv<br>ISM_counts.csv<br>NIC_NNC_counts.csv | Number of reads in each subcategory for FSMs, ISMs, NICs, and NNCs | - - all-tables | Multiple samples file |
| err_counts.csv | Number and percentage of reads with evidence of intrapriming, RT-switching, and non-canonical junctions per samples | - - all-tables | Multiple samples file |
| pca_loadings.csv | PC1 and PC2 loadings from PCA | -pca-tables | Summary file |
| pca_variance.csv | Variance explained by each PC | -pca-tables | Summary file |
| gene_classification.csv | For genes with coverage over a user-defined threshold, gives the annotation category of each gene | Yes | Summary file |
| putative_underannotation.csv | Metrics on NIC and NNC UJCs and flags putative novel transcripts | Yes | Summary file |

The input files for SQANTI-reads include: 1) a GTF file of read alignments, 2) a reference genome FASTA file, 3) a GTF file of the reference transcript model annotation, and 4) a design file containing metadata for multiple samples. The first step of SQANTI-reads involves using the SQANTI3-QC module to generate SQANTI3-like classification and junction files, with the classification file containing one row for each mapped read. Reads are classified according to the SQANTI categories (Tardaguila et al. 2018) as full-splice match (FSM), incomplete-splice match (ISM), novel-in-catalog (NIC), novel-not-in-catalog (NNC), antisense, fusion, genic genomic, and intergenic. SQANTI3 subcategories are also included, based on 5’ and 3’ end positions relative to the annotated transcription start sites (TSS) and transcription termination sites (TTS) (Pardo-Palacios et al. 2024a). Additionally, the reverse transcriptase (RT) switching algorithm of SQANTI3-QC identifies reads with evidence of RT switching events, while reads with more than 60% adenines in the 20 bp downstream of the reported TTS at the genomic level are flagged as potential intrapriming events. The length of each read and the number of exons in each read are also recorded in the classification file.

### Junction Metrics

The SQANTI-reads junction file follows the same format as the SQANTI3 junction file, with each row representing a junction in a read, including the start and end positions of the junction. The distance from the junction start and end to the nearest annotated junction start and end in the reference GTF is calculated. It’s important to note that the nearest annotated start and end positions may not belong to the same annotated junction. SQANTI classifies junctions as known or novel, and as canonical or non-canonical, based on the dinucleotide pairs at the junction’s start and end. By default, dinucleotide combinations of GT-AG, GC-AG, and AT-AC are considered canonical, while any other combinations are classified as non-canonical, although the user can specify additional canonical sites.

SQANTI-reads introduces new metrics to evaluate the relationship between the junctions in mapped reads and the annotated donors and acceptors. In the SQANTI3-QC junction file, the distance from each donor/acceptor in each read to the nearest annotated donor/acceptor is recorded. In SQANTI-reads, the mean absolute distance in nucleotides from the annotated donor/acceptor site, the standard deviation, and the coefficient of variation (CV = standard deviation/mean) are calculated and included in the cv.csv file. Each detected junction is classified as 1) Reference Match junction if the mean distance and the standard deviation to an annotated junction are both equal to 0; 2) CV = 0 junction when the mean distance is greater than 0 and the standard deviation equals 0, and 3) CV > 0 junction when the CV is greater than 0.

### Unique Junction Chain and gene-level information

SQANTI-reads groups mapped reads based on their full junction pattern and refer them to as Unique Junction Chains (UJCs). Each UJC is labeled with a string that includes the chromosome and junction coordinates (Nanni et al. 2024). To enhance computational efficiency, UJC strings are encoded as an index in a hash table (JxnHash). The read count for each JxnHash is calculated and included in the ujc_counts.csv file. Additionally, the number of known canonical, known non-canonical, novel canonical, and novel non-canonical junctions within each UJC is annotated, along with the SQANTI structural category of the UJC. The number of reads within each structural category for each gene, as well as the total number of reads per gene, is stored in the summary file gene_counts.csv.

### Identifying genes that may be under-annotated and transcripts that may be mis-annotated

For expressed genes, a high proportion of reads from a UJC classified as NIC/NNC may indicate the presence of a potentially novel transcript. SQANTI-reads includes a customizable pipeline to identify genes with such potential under-annotation events. The procedure identifies NIC/NNC UJCs that meet a minimum number (R) and proportion (P) of reads, with default values set at 100 reads and 20%, respectively. To mitigate the risk that the NIC/NNC UJC is merely a degradation product, an additional condition is applied: the candidate UJC must include at least 80% of the gene’s junctions (Q) (Figure 3A). The R, P, and Q thresholds are pipeline parameters that can be adjusted by the user. Furthermore, SQANTI-reads allows for the evaluation of under-annotated genes and novel transcripts within a specific subset of samples associated with a particular experimental factor (e.g., developmental stage or technology) using the --factor-level option.

### Multi-sample processing

SQANTI-reads processes multiple samples to generate classification and junction files when a design file (Supplementary file 1) is provided to the sqanti-reads.py command. If individual samples have already been pre-processed with SQANTI3-QC, SQANTI-reads can be run in --fast mode, where the design file links the individual classification and junction files to sample IDs for the calculation of SQANTI-reads metrics, summaries, and a series of visualizations. If pre-processing has not been done, SQANTI-reads is run in --simple mode where SQANTI3 is run on each sample, followed by the calculation of SQANTI3 metrics and summaries. The output also includes a summary for each sample, reporting the mean, median, upper quartile, and lower quartile of mapped read length, as well as the number and proportion of reads that are shorter than 1 kb, between 1 and 2 kb, between 2 and 3 kb, and greater than 3 kb in length, all of which are included in the length_summary.csv file.

### *Drosophila melanogaster* data

A total of 24 samples corresponding to 2 developmental stages (0-1 hours and 3-8 hours post-hatching) and four genotypes (dmel 11037, 11255, 12272 and 12279) (3 samples per experimental condition) were sequenced using Oxford Nanopore Technology (ONT). One barcoded cDNA library was built per sample, the 12 samples from the 0-1 hour and the 12 samples from the 3-8 day stages were pooled and sequenced on a MinION. Data were evaluated with PycoQC (Leger and Leonardi 2019) which focuses on read length and base quality. All samples passed this basic QC, libraries were re-pooled and run on the PromethION. Detailed metadata for these samples are provided in Supplementary Table 1. Raw electrical data were processed with the default Guppy basecaller. Samples were stored in .fast5 format and as .fastq files. The .fast5 files were converted to the Dorado compatible .pod5 format using pod5 (v 0.3.6) and then processed in Dorado (v 0.5.2) (https://github.com/nanoporetech/dorado) using options --recursive --device “cuda:0,1” --kit-name SQK-PCB109 --trim none. Reads were demultiplexed using the demux mode of Dorado (v 0.5.2) with options --no-classify --emit-fastq resulting in separate Dorado fastq files for each sample. The fastq files generated by Guppy/Dorado were both processed using pychopper (v 2.7.1), the oriented fastq files were aligned to *D. melanogaster* 6.50 and the resulting sam files were converted to gtf using samtools (v 1.10) (Li et al. 2009) and bedtools (v 2.29.2) (Quinlan and Hall 2010). The resulting gtf files (67 technical replicates from 24 samples), the *D. melanogaster* 6.50 fasta reference file (https://ftp.flybase.net/releases/FB2023_01/dmel_r6.50/ (Öztürk-Çolak et al. 2024)), and a design file (Supplementary File 2) were used as input to SQANTI-reads. The *Drosophila* dataset includes, therefore, two experimental conditions (time and genotype) and two technical conditions (sequencing platform and base caller) with the experimental samples multiplexed and evaluated with both technical conditions. The SQANTI-reads output for this dataset is provided in Supplementary File 3.

### Human Cell line WTC11

We used publicly available lrRNA-seq data from the Long-read RNA-seq Genome Annotation Assessment Project (Pardo-Palacios et al. 2024b) to illustrate the utility of SQANTI-reads. Specifically, we used triplicate measurements of the transcriptome of the WTC11 human cell line that were profiled by cDNA PacBio Sequel II, cDNA Oxford Nanopore Minion, and direct RNA Oxford Nanopore Minion methods. Data were downloaded from the ENCODE website (https://www.encodeproject.org/search/?type=Experiment&internal_tags=LRGASP). Accession numbers for these samples are provided in Supplementary Table 2. The fastq files were pre-processed by LRGASP researchers as described in (Pardo-Palacios et al. 2024b). We used the gtf files of read alignments, GENCODE’s GRCh38.p13 reference genome gtf and fasta for release 38 (https://www.gencodegenes.org/human/release_38.html), and a design file (Supplementary file 4) to run SQANTI-reads on the WTC11 samples. The SQANTI-reads output is provided in Supplementary File 5.

## Acknowledgements

This work was supported in part by a grant from the National Institutes of Health (1R21HG011280-01), the Spanish MICIN (PID2020-119537RB-I00), the European Union’s programme Horizon Europe under the Marie Skłodowska-Curie Actions postdoctoral fellowship to C.M. (101149931). This work is also supported in part by a grant from the National Cancer Institute (NCI P01 CA214091), the National Institute of General Medical Sciences (NIGMS GM137430), the University of Florida Department of Molecular Genetics and Microbiology, the University of Florida Genetics Institute, the University of Florida Cancer Center, and the University of Florida Research Computing Center (www.rc.ufl.edu) and the Latin American and Caribbean Scholars award to N.K. Part of the computations were performed on the high performance computing cluster Garnatxa at the Institute for Integrative Systems Biology (I2SysBio). I2SysBio is a joint research center formed by University of Valencia (UV) and Spanish National Research Council (CSIC). We acknowledge Ashley Myrick for help with some of the initial CV plots and Knife Bankole for the initial coding of the junction hash. Alison Morse prepared all samples and libraries for the *Drosophila* experiment, ran all initial QC analyses and recalled all the bases for ONT data. We acknowledge Rolf Renne for his support.

## Author contributions

N.K. conducted research, developed software implementations, created figures and drafted manuscript. C.M. revised code, integrated new methods into SQANTI Github and helped with data analysis. L.M. conceived and supervised the study and drafted the manuscript. A.C. supervised the study, contributed to conceptualizations and drafted the manuscript.

## Data access

*Drosophila* long read data are available at the sequencing read archive (SRA) under the Bioproject PRJNA1134728.

---

**Supplementary Figure 1:**
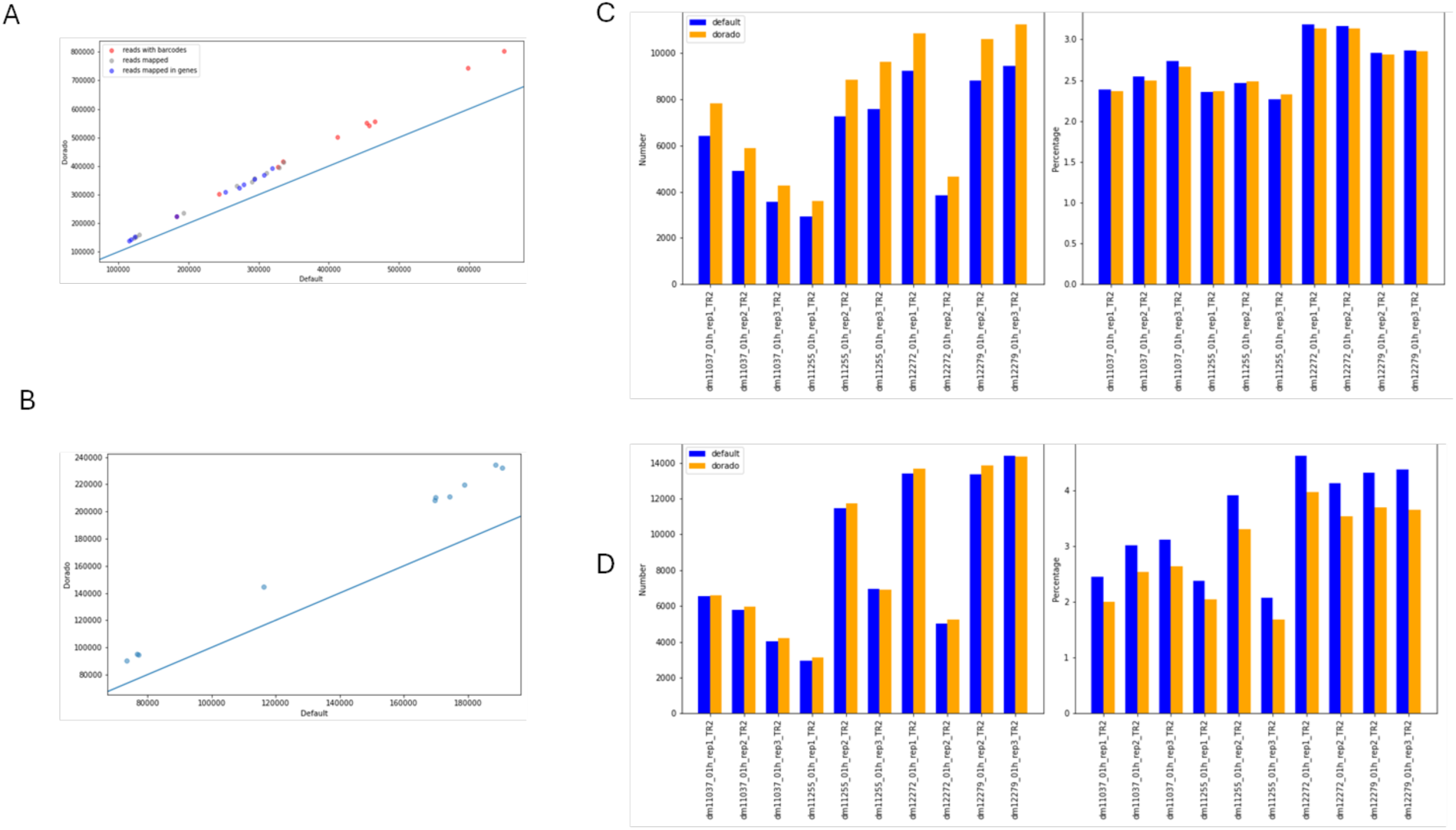
Comparison of Guppy and Dorado basecalling for technical replicate 2. A) Number of reads with assignable barcodes (red), number of mapped reads (grey) and number of reads mapped in genes (blue) in Dorado vs Guppy. B) Number of reads >2kb in Dorado vs Guppy C) Number and percentage of reads with evidence of intrapriming in Dorado and Guppy D) Number and percentage of reads with non-canonical junctions in Dorado and Guppy.

**Supplementary Figure 2:**
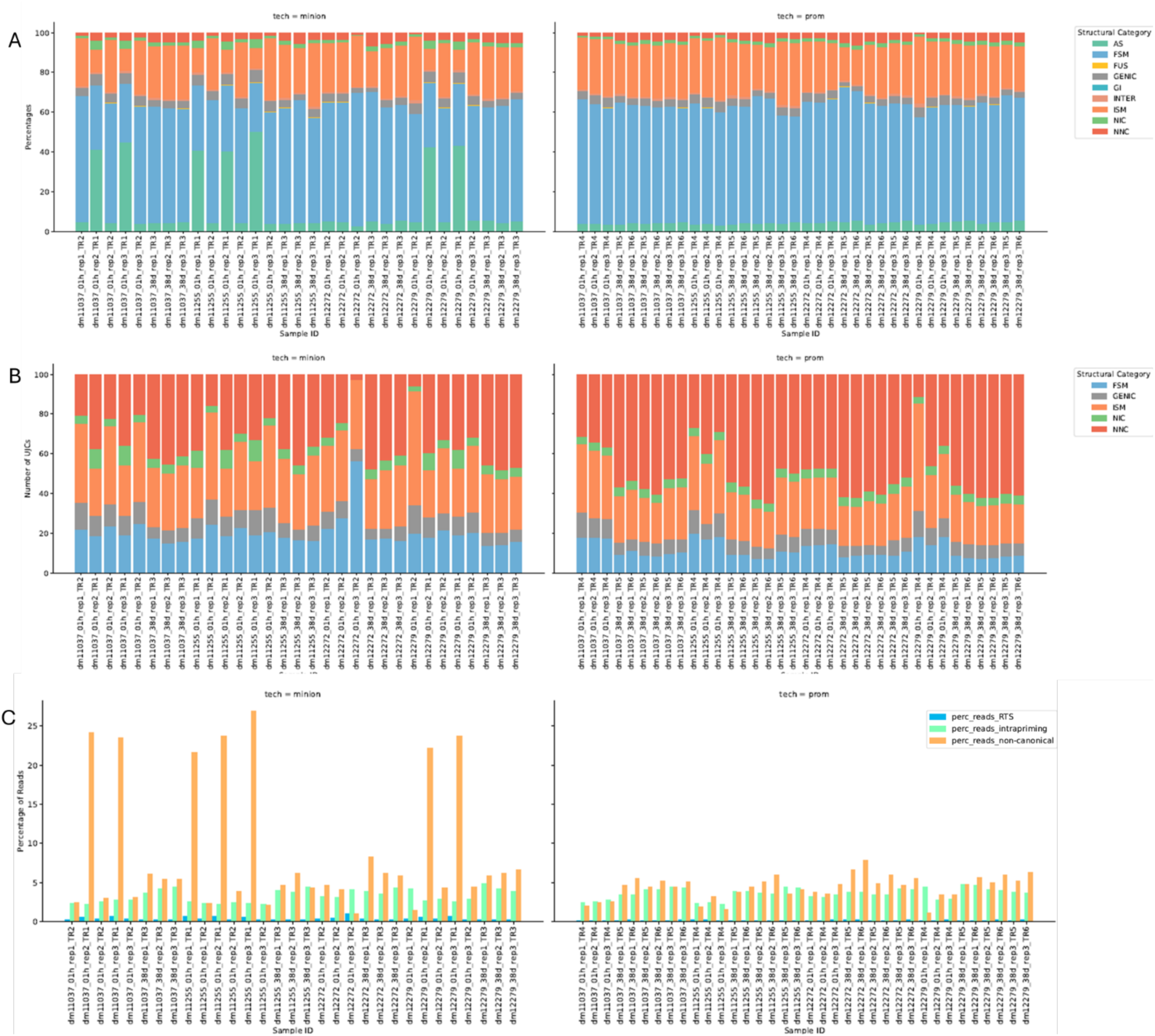
*Drosophila* technical replicates basecalled with Dorado. A) Proportion of reads in each structural category. B) Proportion of unique junction chains (UJCs) in each structural category. C) Proportion of reads with evidence of RT-switching (blue), intrapriming (green) and non-canonical junctions (orange)

**Supplementary Figure 3:**
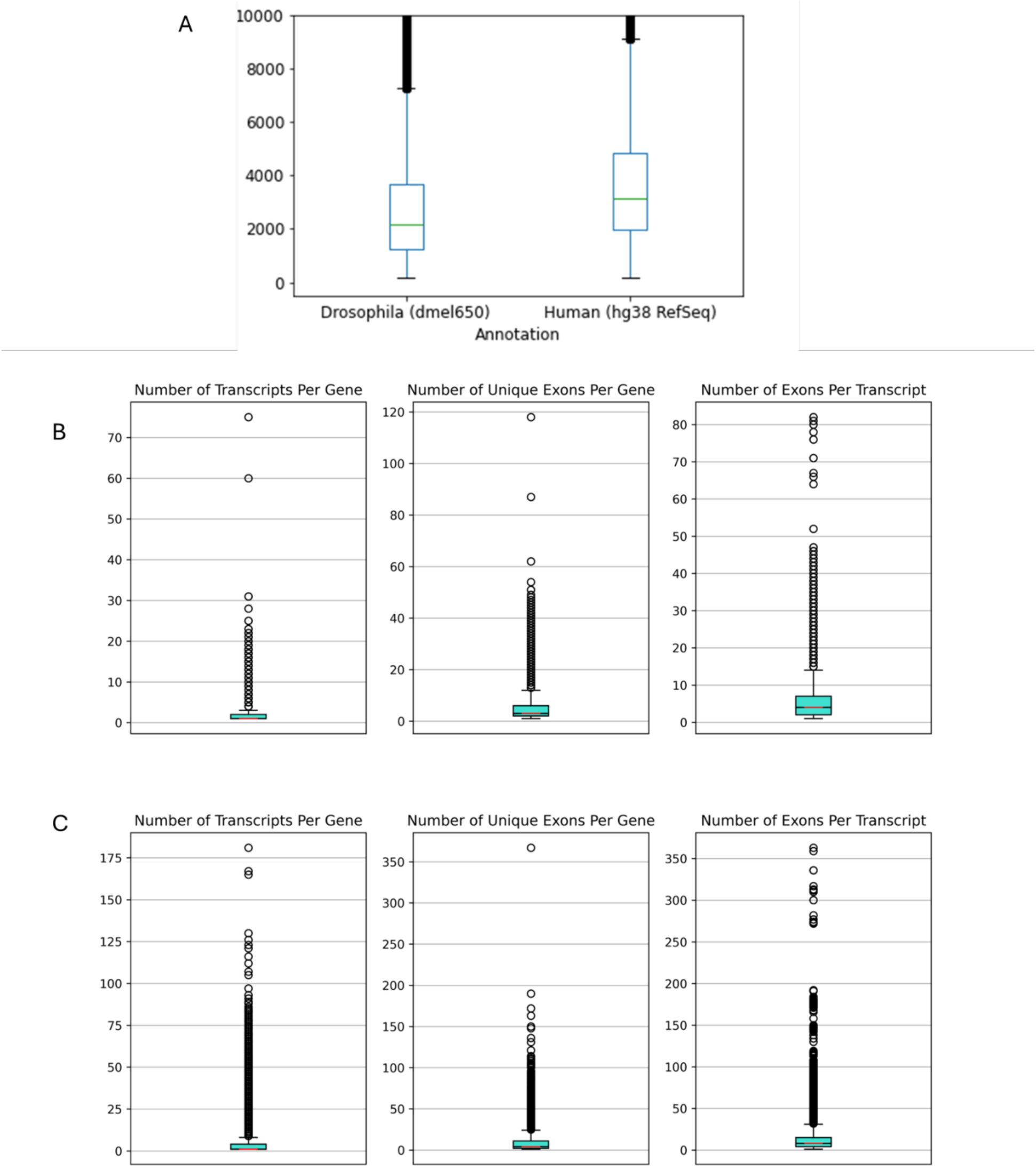
Complexity Metrics of Drosophila (dmel650) and Human (hg38 Refseq) Annotations. A) Distribution of protein-coding transcript lengths in dmel650 and hg38 Refseq. B) Transcriptome Complexity Metrics for dmel650 C) Transcriptome complexity metrics for hg38 Refseq

**Supplementary Figure 4:**
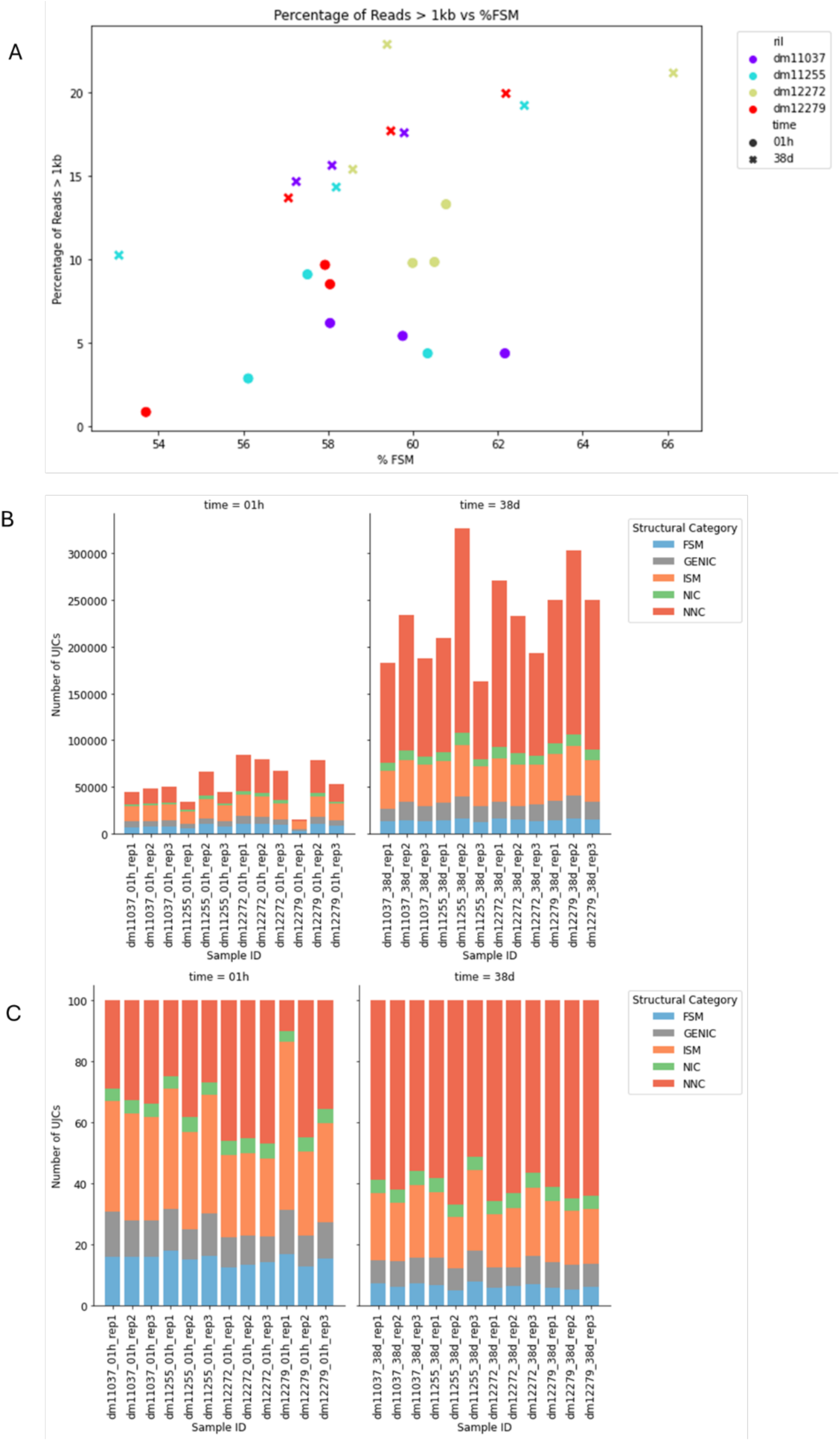
SQANTI-reads plots for *Drosophila* samples basecalled with Dorado A) Percentage of reads greater than 1kb vs Percentage of reads classified as full-splice match (FSM) B) Number of UJCs detected coloured by structural category of the UJC. C) Proportion of UJCs coloured by structural category of the UJC

**Supplementary Figure 5:**
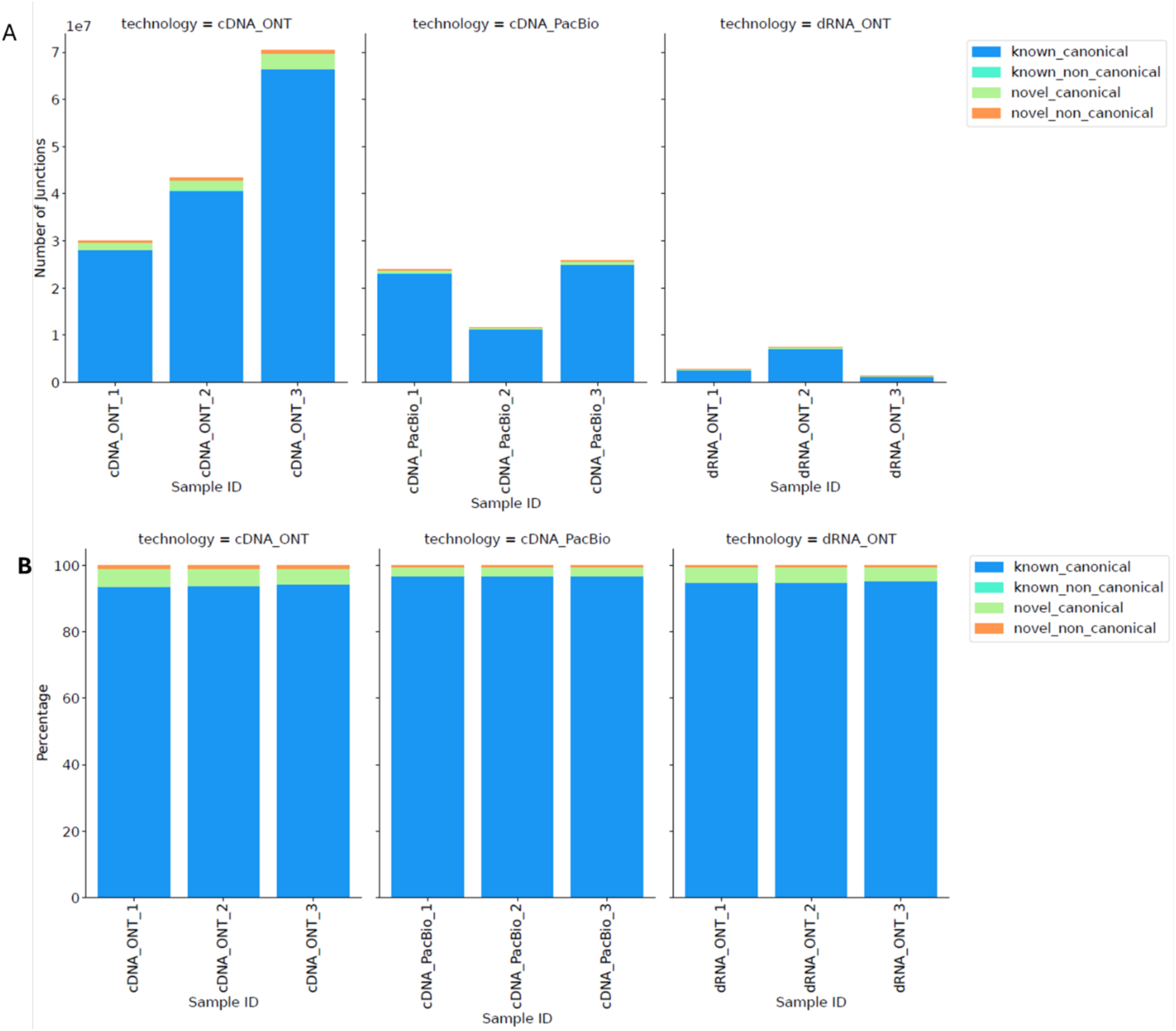
Junction plots for WTC11 samples A) Number of junctions in reads coloured by junction categories (known-canonical, known-non-canonical, novel-canonical and novel-non-canonical) B) Proportion of junctions in reads in each junction category

**Supplementary Figure 6:**
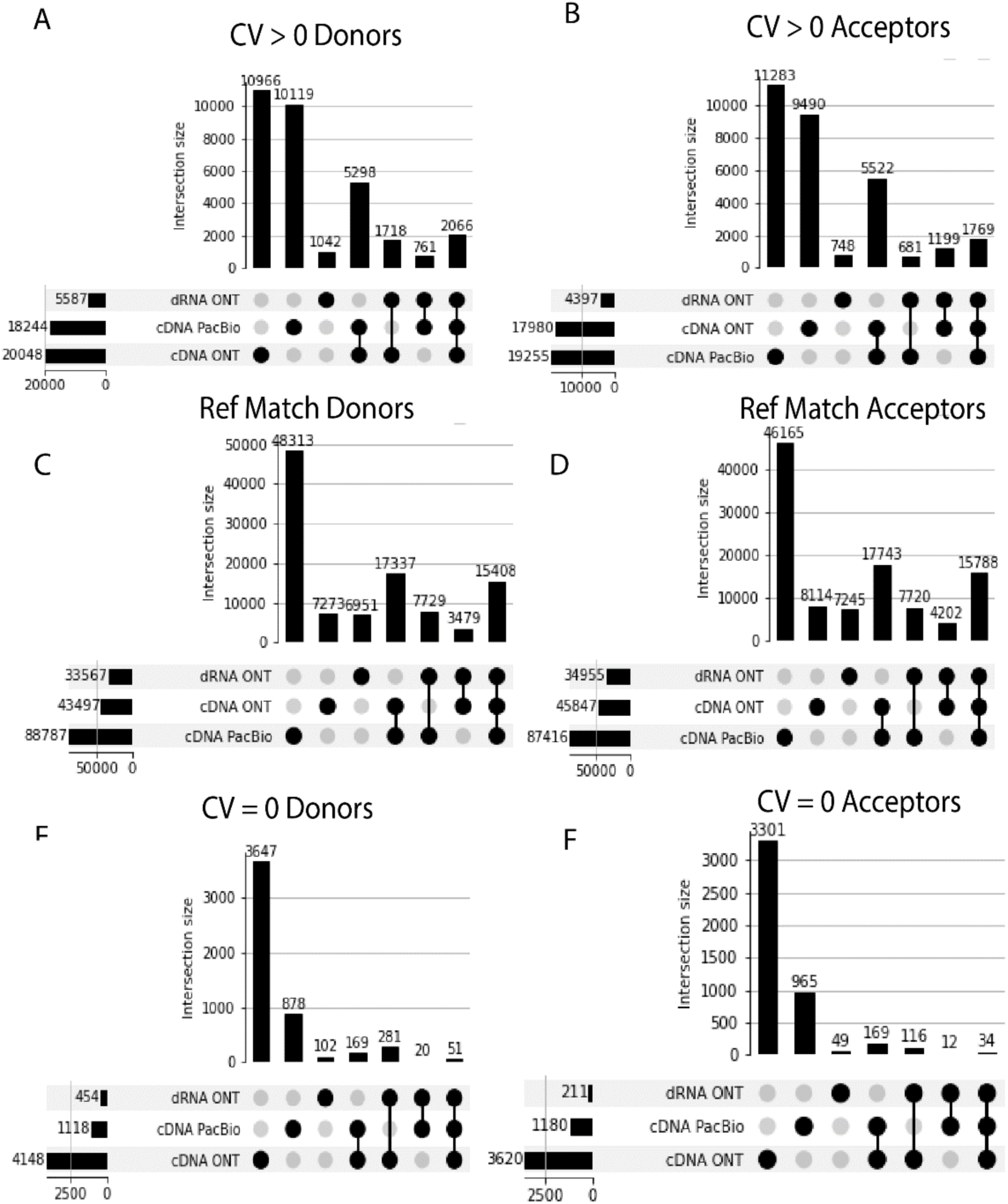
Upset plot of donors and acceptors detected in replicate 1 of the WTC11 samples of each technology by variation category. CV > 0 donors (A) and acceptors (B). Reference match donors (C) and acceptors (D). CV = 0 donors (E) and acceptors (F)

**Supplementary Figure 7:**
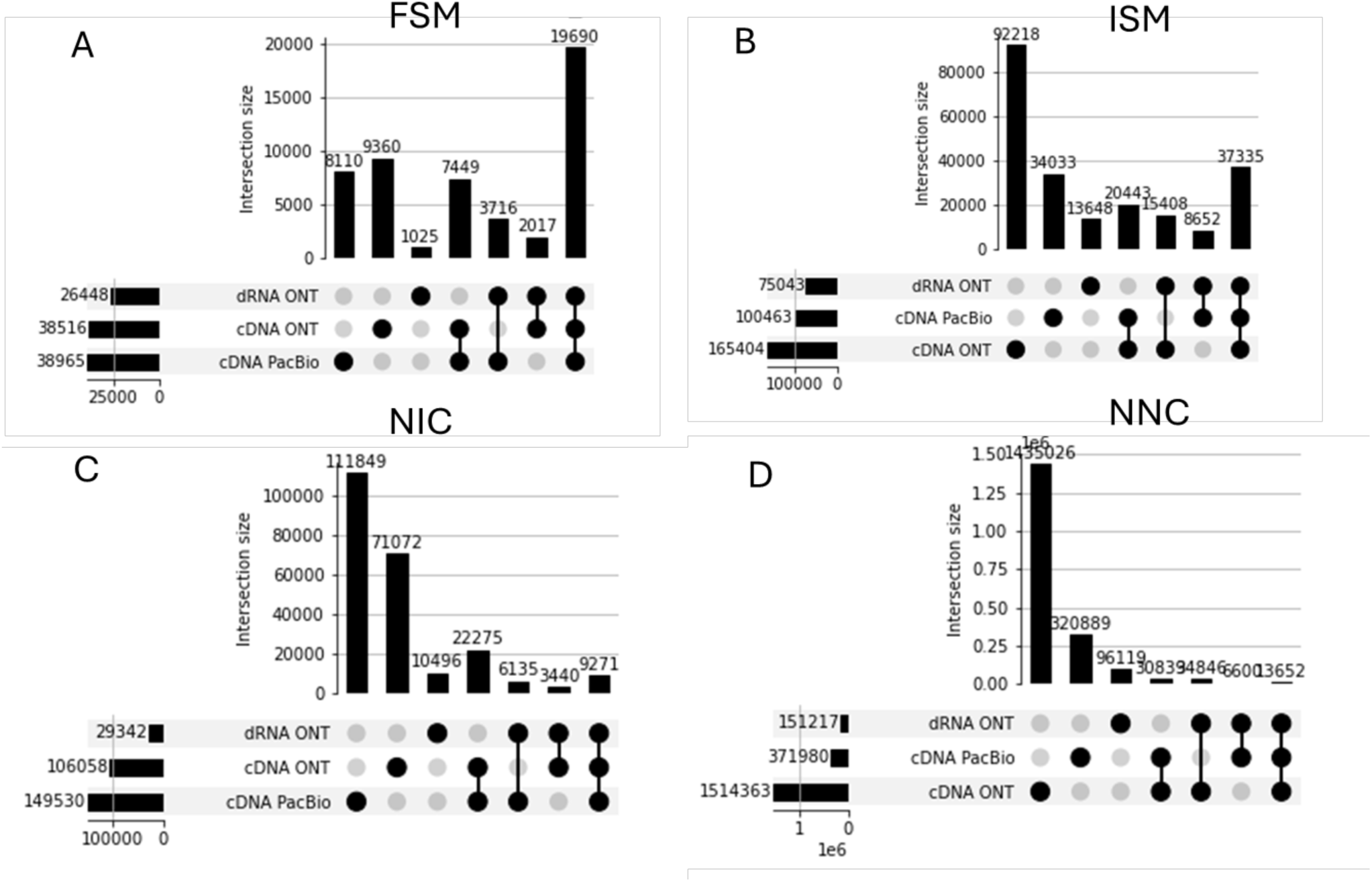
Upset plot of multi-exon UJCs detected in each technology by structural category in the WTC11 samples. A) FSM UJCs B) ISM UJCs C) NIC UJCs and D) NNC UJCs

**Supplementary Figure 8:**
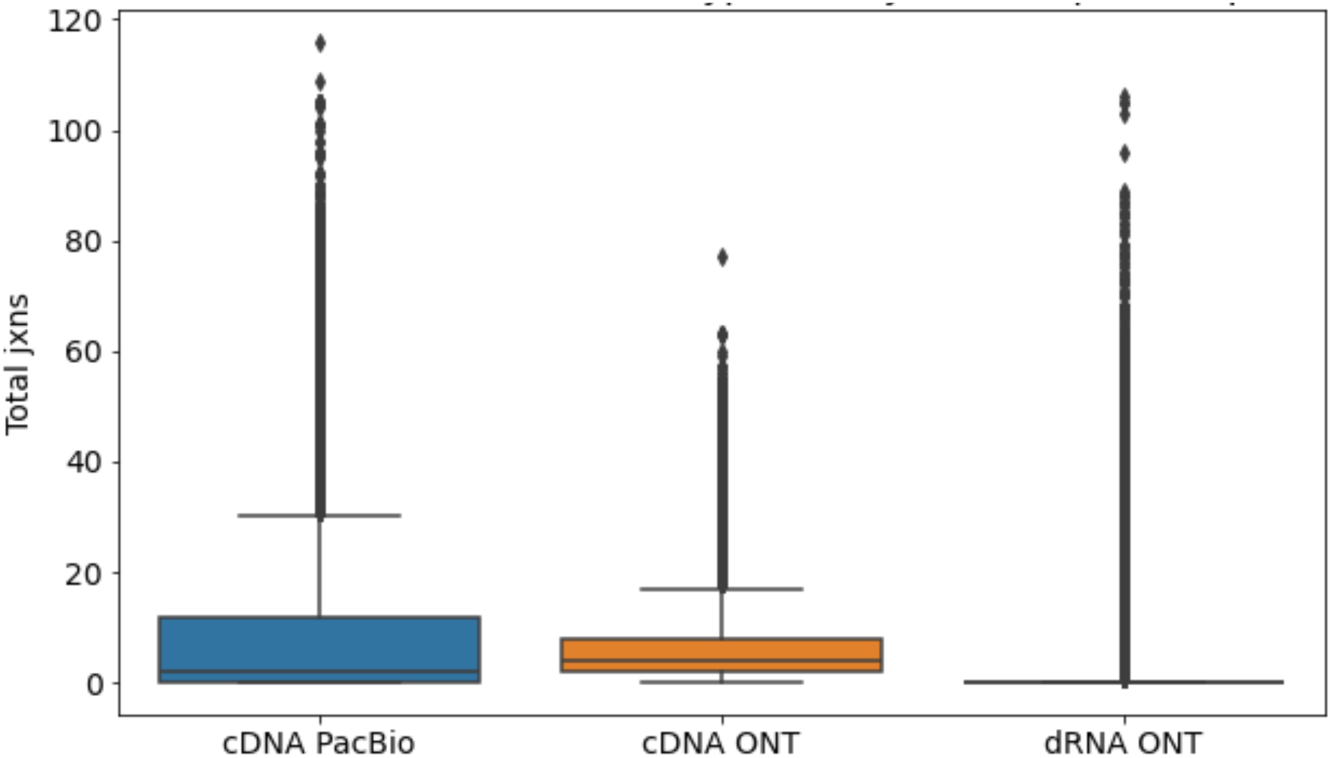
Distribution of number of junctions in FSM UJCs detected exclusively in one technology for the WTC11 samples

**Supplementary Figure 9:**
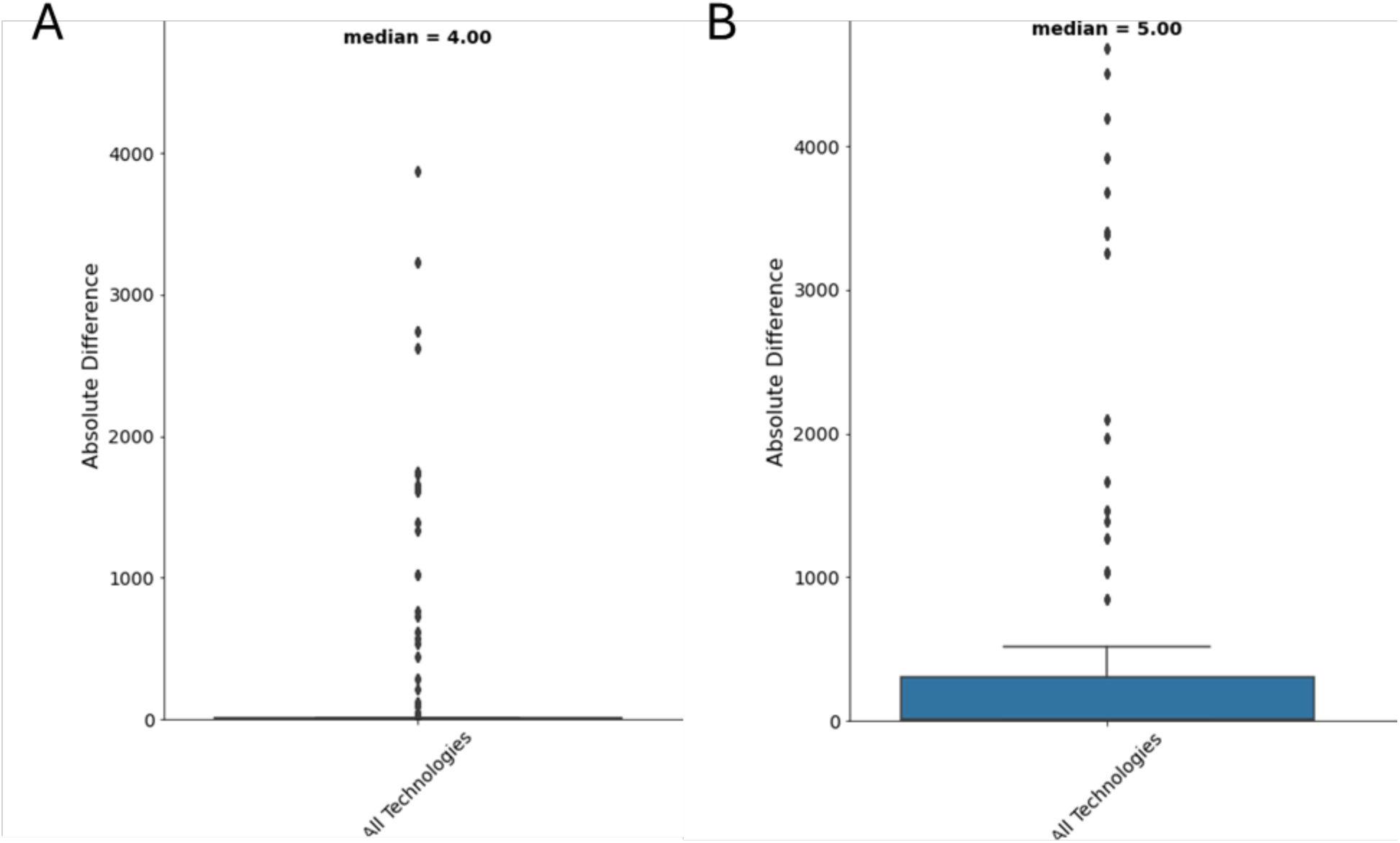
For donors (A) and acceptors (B) with cv=0 in replicate 1 of the WTC11 samples, boxplot of the distribution of the distance of the read donors and acceptors to the annotated donors and acceptors in replicate 1.

**Supplementary Figure 10:**
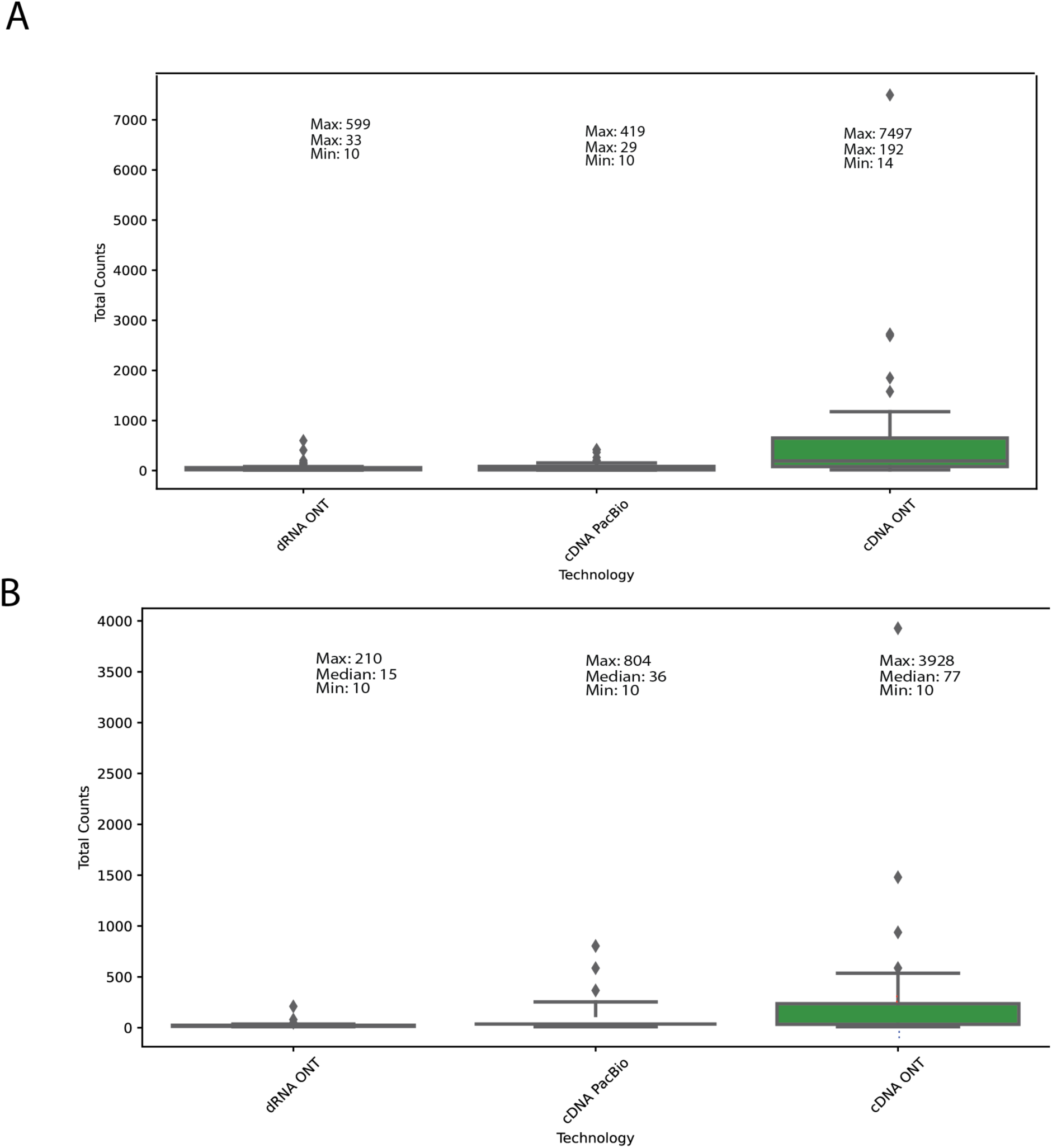
For donors (A) and acceptors (B) with cv=0 in replicate 1 of the WTC11 samples, boxplot of the number of reads associated with the reference junction with cv=0.

**Supplementary Figure 11:**
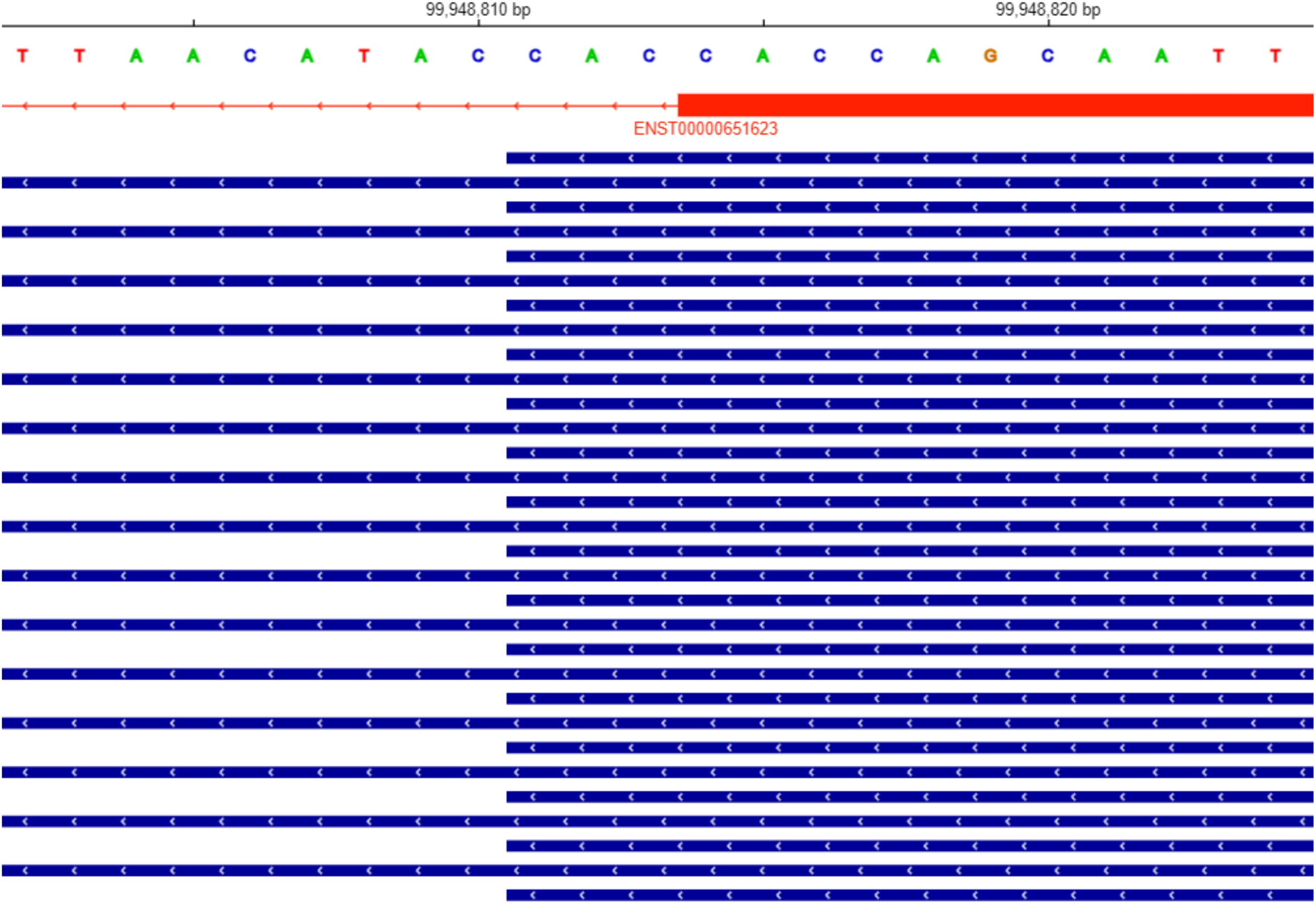
IGV visualization of a reference donor with CV = 0 in the H2AZ1 gene at position 99,948,814 on chromosome 4. All of the reads associated with this donor map to position 99,948,811 which is 3 nts away from the annotated donor position. The annotated transcript is shown in red and the reads are shown in blue.

**Supplementary Table 1:** Metadata for Drosophila Experiment.

| Sample | Genotype | Stage | Number of Technical Replicates | Minion Runs | Promethion Runs |
| --- | --- | --- | --- | --- | --- |
| dm11037_01h_rep1 | 11037 | 01h | 2 | 1 | 1 |
| dm11037_01h_rep2 | 11037 | 01h | 3 | 2 | 1 |
| dm11037_01h_rep3 | 11037 | 01h | 3 | 2 | 1 |
| dm11255_01h_rep1 | 11255 | 01h | 3 | 2 | 1 |
| dm11255_01h_rep2 | 11255 | 01h | 3 | 2 | 1 |
| dm11255_01h_rep3 | 11255 | 01h | 3 | 2 | 1 |
| dm12272_01h_rep1 | 12272 | 01h | 2 | 1 | 1 |
| dm12272_01h_rep2 | 12272 | 01h | 2 | 1 | 1 |
| dm12272_01h_rep3 | 12272 | 01h | 2 | 1 | 1 |
| dm12279_01h_rep1 | 12279 | 01h | 2 | 1 | 1 |
| dm12279_01h_rep2 | 12279 | 01h | 3 | 2 | 1 |
| dm12279_01h_rep3 | 12279 | 01h | 3 | 2 | 1 |
| dm11037_38d_rep1 | 11037 | 38d | 3 | 1 | 2 |
| dm11037_38d_rep2 | 11037 | 38d | 3 | 1 | 2 |
| dm11037_38d_rep3 | 11037 | 38d | 3 | 1 | 2 |
| dm11255_38d_rep1 | 11255 | 38d | 3 | 1 | 2 |
| dm11255_38d_rep2 | 11255 | 38d | 3 | 1 | 2 |
| dm11255_38d_rep3 | 11255 | 38d | 3 | 1 | 2 |
| dm12272_38d_rep1 | 12272 | 38d | 3 | 1 | 2 |
| dm12272_38d_rep2 | 12272 | 38d | 3 | 1 | 2 |
| dm12272_38d_rep3 | 12272 | 38d | 3 | 1 | 2 |
| dm12279_38d_rep1 | 12279 | 38d | 3 | 1 | 2 |
| dm12279_38d_rep2 | 12279 | 38d | 3 | 1 | 2 |
| dm12279_38d_rep3 | 12279 | 38d | 3 | 1 | 2 |

**Supplementary Table 2:** Metadata for WTC11 experiment.

| Sample | Library Prep | Platform | Run Accession | Rep | Biosample | File Accession |
| --- | --- | --- | --- | --- | --- | --- |
| WTC11 | dRNA | ONT | ENCSR392BGY | 1 | ENCBS944CBA | ENCFF155CFF |
| WTC11 | dRNA | ONT | ENCSR392BGY | 2 | ENCBS593PKA | ENCFF771DIX |
| WTC11 | dRNA | ONT | ENCSR392BGY | 3 | ENCBS474NOC | ENCFF600LIU |
| WTC11 | cDNA | PacBio | ENCSR507JOF | 1 | ENCBS944CBA | ENCFF563QZR |
| WTC11 | cDNA | PacBio | ENCSR507JOF | 2 | ENCBS593PKA | ENCFF370NFS |
| WTC11 | cDNA | PacBio | ENCSR507JOF | 3 | ENCBS474NOC | ENCFF245IPA |
| WTC11 | cDNA | ONT | ENCSR539ZXJ | 1 | ENCBS944CBA | ENCFF263YFG |
| WTC11 | cDNA | ONT | ENCSR539ZXJ | 2 | ENCBS593PKA | ENCFF023EXJ |
| WTC11 | cDNA | ONT | ENCSR539ZXJ | 3 | ENCBS474NOC | ENCFF961HLO |

